# Meta-transcriptomic discovery of a divergent circovirus and a chaphamaparvovirus in captive reptiles with proliferative respiratory syndrome

**DOI:** 10.1101/2020.08.24.264143

**Authors:** Wei-Shan Chang, Ci-Xiu Li, Jane Hall, John-Sebastian Eden, Timothy H. Hyndman, Edward C. Holmes, Karrie Rose

**Author notes:** Correspondence: Edward Holmes - Karrie Rose.

## Abstract

Viral pathogens are being increasingly described in association with mass morbidity and mortality events in reptiles. However, our knowledge of reptile viruses and their role in population health remains limited. Herein, we describe a meta-transcriptomic investigation of a mass morbidity and mortality event in a colony of central bearded dragons (*Pogona vitticeps*) in 2014. Severe, extensive proliferation of the respiratory epithelium was consistently found in affected dragons. Similar proliferative lung lesions were identified in bearded dragons from the same colony in 2020 in association with increased intermittent mortality. Total RNA sequencing of bearded dragon tissue identified two divergent DNA viruses: a reptile-infecting circovirus, denoted bearded dragon circovirus (BDCV), and the first exogeneous reptilian chaphamaparvovirus - bearded dragon chaphamaparvovirus (BDchPV). Phylogenetic analysis revealed that BDCV was most closely related to bat-associated circoviruses, exhibiting 70% amino acid sequence identity. In contrast, the newly discovered BDchPV showed approximately 35-40% identity in the non-structural (NS) protein to parvoviruses obtained from tilapia fish and crocodiles in China. Subsequent specific PCR assays detected BDCV exclusively and comprehensively within animals with proliferative pulmonary lesions and respiratory disease. This study expands our understanding of viral diversity in the context of diseased reptiles in captivity.

## Introduction

Squamates (scaled reptiles) are one of the most diverse groups of vertebrate fauna in Australia [1]. Bearded dragons (*Pogona spp*.), including central bearded dragons (*Pogona vitticeps*), are native reptiles that inhabit wide geographic areas throughout the arid and semi-arid regions of Australia. Bearded dragons are increasingly popular exotic pets, and are now one of the most common companion lizard species in Australia, Europe, Asia, and North America [2]. Bearded dragons are also ideal for laboratory studies: their well-established history in captivity makes them suitable for investigations of reptile physiology, and they have unusual patterns of chromosomal, genomic and temperature-based sex-determination, including sex reversal with specific behavioral consequences [3,4]. Despite their significance, our knowledge of reptile pathogens, particularly viruses, remains limited. Indeed, to date the only viruses known to infect bearded dragons are double-strand DNA viruses from the families *Iridoviridae* [5–8] and *Adenoviridae* [9,10], single-stranded DNA viruses from the *Parvoviridae* [11–13] and RNA viruses from the *Paramyxoviridae* [14,15].

The diagnosis of microbial infections in reptiles is complex and often associated with multiple, synergistic and predisposing factors. It is likely that undetected viruses contribute to a number of infectious disease outbreaks in captive reptiles, including bearded dragons. To this end, we investigated two unusual mortality events of bearded dragons in a research colony in Australia in 2014 and 2020. In 2014, approximately 40 central bearded dragons died after emergence from brumation in a colony of over 400 animals located in Canberra, Australia. Affected animals were either found dead or were found listless and died within 24 hours. In 2020, the same colony experienced low grade post-brumation mortality followed several months later by star gazing and poor body condition in juveniles and acute death in an adult bearded dragon.

Brumation is an extreme hypometabolic state used by some reptiles to cope with low or unpredictable food availability and unfavorable seasonal conditions for the duration of winter. A number of specific protective measures have been identified during this hibernation in bearded dragons, including increased neuroprotection in the brain, maintenance of heart function through hypertrophy, and upgrading antioxidant capacity and mitochondrial maintenance by skeletal muscle atrophy [16]. Emergence from brumation may also represent a period of increased disease susceptibility in reptiles, as shown by downregulated transcription of several genes responsible for microbial pathogen defense, cellular and oxidative stress, and cell differentiation and growth [17].

Herein, we employed PCR testing and meta-transcriptomic approaches combined with gross and microscopic pathology to investigate the possible involvement of viral pathogens associated with unknown mortality events in a captive colony of bearded dragons.

## Materials and Methods

### Sample collection and processing

Tissues were collected from two disease outbreaks in a research colony in Canberra, Australia. Five dead bearded dragons were submitted for post-mortem examination during a mortality event in 2014. In 2020, 12 live, sick animals were examined, sedated with 20 mg/kg of im alfaxalone (Alfaxan® CD-RTU 10mg/mL, Jurox, Australia), delivered into the biceps to rapidly induce a surgical plane of anesthesia, followed by 1ml/kg of iv pentobarbitone (Lethobarb, 325mg/mL, Virbac, Australia) to effect euthanasia. Gross post-mortem examinations were conducted immediately, and fresh portions of cervical spinal cord, liver, heart, spleen and kidney were collected aseptically and frozen at −80°C. A range of tissues were fixed in 10% neutral buffered formalin, processed in ethanol, embedded with paraffin, sectioned, stained with hematoxylin and eosin, and mounted with a cover slip prior to examination by light microscopy. Giemsa, Ziehl-Neelsen and periodic acid-Schiff stains were applied to a subset of embedded tissue samples to identify bacteria, and to exclude the presence of mycobacteria and fungi. Aerobic, anaerobic and fungal cultures were undertaken on lung and liver samples aseptically collected from the five dragons that died in 2014.

Proliferation of respiratory epithelium was graded on a scale of 0-4, where 0 indicated no discernable lesion and 4 represented severe and extensive proliferation of infundibular and mesobronchial epithelium. Pulmonary and hepatic inflammation were graded on a scale of 0-4, where 0 indicated no discernible lesions, 1 represented scattered leukocytes and 4 represented severe and extensive heterophilic inflammation. Pulmonary and hepatic necrosis were graded on a scale of 0-4 where 0 indicated no discernible necrosis, 1 represented mild single cell degeneration or necrosis, and 4 characterized extensive caseating necrosis. Samples were collected under the Opportunistic Sample Collection Program of the Taronga Animal Ethics Committee. Samples were submitted to Murdoch University for PCR testing of pneumotropic viruses, and to the University of Sydney for meta-transcriptome analysis and further PCR testing.

For PCR testing for pneumotropic viruses, total nucleic acid was extracted from tissues using the MELT™ Total Nucleic Acid Isolation System (Ambion, USA) according to the manufacturer’s instructions. Total nucleic acid was eluted into 30 μL of elution buffer. For meta-transcriptomics and PCR testing for parvoviruses and circoviruses, the RNA extracted from the spinal cord, liver, lung and kidney of diseased animals was processed using the RNeasy Plus Mini Kit (Qiagen, Germany). Initially, frozen tissue was partially thawed and submerged in RLT lysis buffer containing 1% ß-mercaptoethanol and 0.5% reagent DX before being homogenized using a hand-operated TissueRupture (Qiagen). To pool samples in equal proportions, RNA concentrations and integrity were validated using a NanoDrop spectrophotometer (ThermoFisher Scientific, USA) and a TapeStation (Agilent, USA). Illumina TruSeq stranded RNA libraries were prepared on the pooled samples following rRNA depletion using a RiboZero Gold rRNA removal kit (Epidemiology). Finally, 150 bp paired-end sequencing of the rRNA-depleted RNA library was generated on an Illumina NovaSeq 6000 service system at Australian Genome Research Facility (AGRF), Melbourne.

### PCR testing for pneumotropic viruses

Nucleic acid from the kidney, liver and lung was pooled and tested for sunshineviruses, orthoreoviruses and ferlaviruses using one-step reverse transcription (RT)-PCR. For each virus genus, 1 μL of extracted nucleic acid was added to 0.8 μL of SuperScript® III RT/Platinum® Taq Mix (Invitrogen, Australia), 10 μL of 2x Reaction Mix, 1 μM (final concentration) of each primer and was then made up to a final volume of 20 μL. For nested PCRs, 1 μL of PCR product was used as template for the second round of amplification using Platinum® PCR Supermix (Invitrogen) in a final volume of 20 μL. For sunshineviruses, the primer pair SunshineS2-SunshineAS2 was used and cycling conditions were 45°C × 45 m, 94°C x_2 m, 40 x_(94°C × 20 s, 51°C × 30 s, 72°C × 20 s) [18]. For orthoreoviruses, the primer pairs 1607F-2608R and 2090F-2334R were used and cycling conditions were 45°C × 45 m, 94°C × 2 m, 40 × (94°C × 20 s, 46°C × 60 s, 72°C × 60 s) and 94°C × 2 m, 40 × (94°C × 20 s, 47°C × 45 s, 72°C × 45 s) for the first and second rounds, respectively [19]. For ferlaviruses, the primer pairs L5-L6 and L7-L8 were used and cycling conditions were 45°C × 45 m, 94°C × 2 m, 40 × (94°C × 20s, 46°C × 60 s, 72°C × 60 s) and 94°C × 2 m, 40 × (94°C × 20 s, 47°C × 45 s, 72°C × 45 s), respectively [20]. Sunshine Coast virus [21], a reptile orthoreovirus (kindly provided by Dr Rachel Marschang) and a ferlavirus (American Type Culture Collection VR-1408) were used as positive controls. PCR products were visualised with agarose gel electrophoresis.

### Pathogen discovery using metatranscriptomics

Our meta-transcriptomic approach to pathogen discovery was based on those previously employed by our group [22,23]. RNA sequencing reads were trimmed for quality using Trimmomatic [24] before *de novo* assembly with Trinity, version 2.5.1 [25]. Assembled sequence contigs were then annotated using both nucleotide BLAST searches against the NCBI nt database and Diamond Blastx against the NCBI nr database [26], with e-value cut-offs of <10^−6^ and <10^−5^, respectively. Open reading frames were predicted from the potential viral contigs using Geneious v11.1.5 [27], with gene annotation and functional predictions made against the Conserved Domain Database (CDD) [28]. To evaluate the virus abundance and coverage, reads were mapped back to the genome with BBmap [29].

### PCR testing for chaphamaparvoviruses and circoviruses

Virus-specific PCR primers were designed based on identified transcripts from the RNA-seq data (Table 1). Accordingly, the extracted RNA (5 μL) from each tissue of individual cases was reverse transcribed using the SuperScript IV VILO cDNA synthesis system (Invitrogen). The cDNA generated from each sampled tissue (2 μL) was used for viral specific PCRs targeting regions identified by RNA-Seq. For BDCV, the primer pair BDCV_F2 and BDCV_R1 were used (cycling conditions = 98°C × 1 m, 35 × (98°C × 10s, 62°C × 10 s, 72°C × 45 s). For BDchPV, the primers BDChPV_F4 and BDChPV_R3 were used (cycling conditions 98°C × 1 m, 35 × (98°C × 10s, 62.2°C × 10 s, 72°C × 60 s). All PCRs were performed using Platinum SuperFi DNA polymerase (Invitrogen) with a final concentration of 0.2 μM for both forward and reverse primers. PCR products were visualized with agarose gel electrophoresis and Sanger sequenced at the AGRF.

**Table 1.**
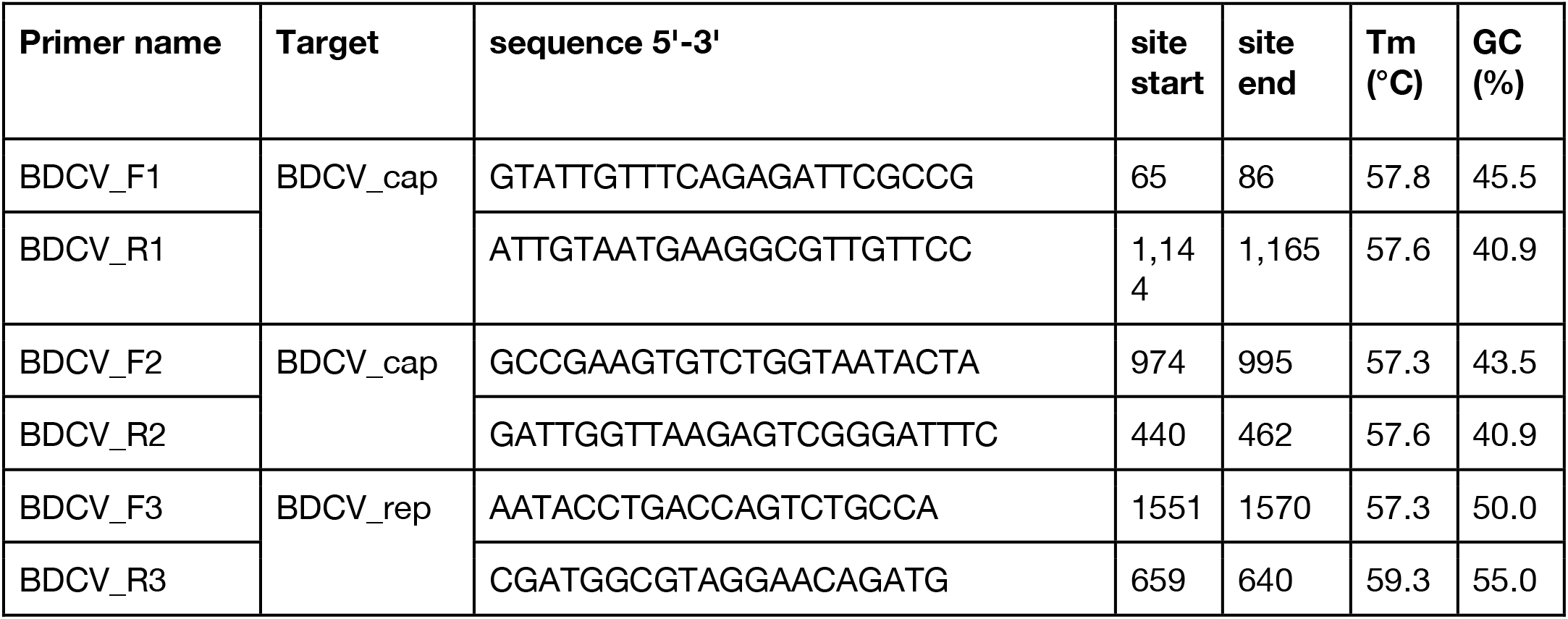
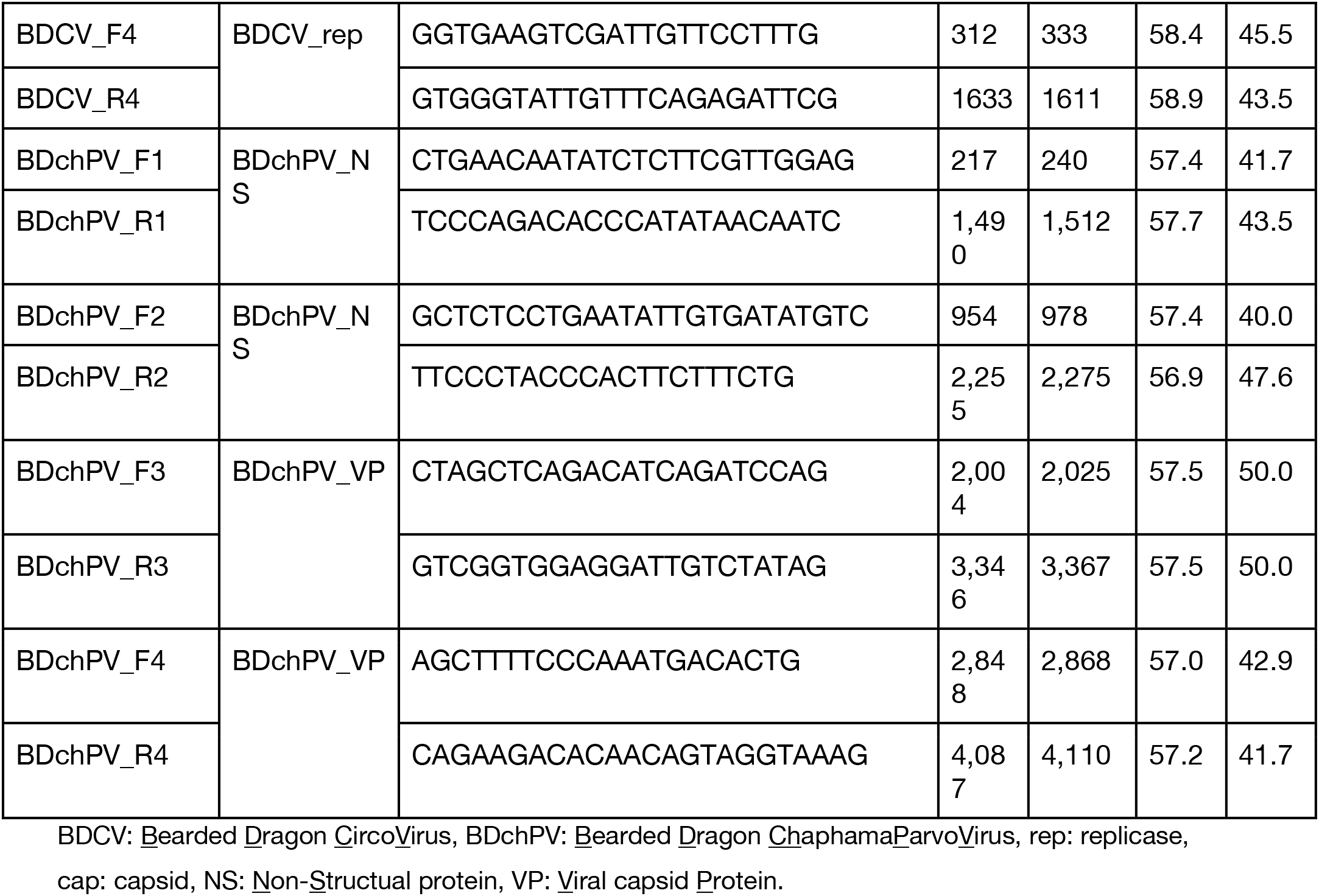
PCR primers used in this study.

### Phylogenetic analysis

Phylogenetic trees were estimated based on MAFFT [30] alignments of the conserved polymerase region of the viruses identified here, with all analyses utilizing representative members of each virus family taken from NCBI/GenBank. Maximum likelihood (ML) phylogenies were inferred using IQ-TREE (version 1.4.3)[31], employing the LG+F+ 4 model of amino acid substitution on the NS protein (717 amino acids) and the LG+F+I+ 4 model on the replicase (rep) protein (274 amino acids). Statistical support for individual nodes was estimated via bootstrap resampling (1000 replicates). Data was visualized using Figtree 1.4.2 (http://tree.bio.ed.ac.uk/software/figtree/).

### Mining the Sequence Read Archive (SRA)

To help determine if BDCV and BDchPV were present in other bearded dragon or other reptile species, we screened publicly accessible high-throughput sequencing data available on the NCBI SRA database (https://www.ncbi.nlm.nih.gov/sra). Accordingly, a large RNA-seq data set was obtained using the NCBI SRA toolkit (version 2.9.2) from the bearded dragon genus *Pogona* (NCBI taxid:52201). Retrieved FASTQ reads were then subjected to a blast analysis using Diamond v.0.9.25 [32] against customized databases containing the core genes from reference circoviruses and parvoviruses, employing an e-value of cut-off of 1 × e^−5^. No additional novel reptilian associated circoviruses and parvoviruses were detected using this approach.

## Results

### Clinical and histopathological findings of diseased captive reptiles

The mass mortality and morbidity of up to 40 adult central bearded dragons in 2014, and intermittent ill health and star gazing in hatchlings from the same research colony in 2020, were both accompanied by unusual and severe proliferation of pulmonary epithelium. The infundibular mucosa, which is normally squamous, exhibited metaplasia ranging from columnar to pseudostratified columnar and ciliate. The mesobronchial epithelium was similarly thickened, pseudostratified, and dysplastic with multifocal loss of cilia. Amorphous basophilic inclusions were evident within enlarged nuclei of infundibular epithelial cells in 2/7 affected animals. Eosinophilic intracytoplasmic inclusion bodies were evident within the respiratory epithelium of 3/7 affected animals. Rarely binucleate epithelial cells and syncytia were identified within mesobronchial epithelium. Similar inclusions were evident within hepatocytes of 3/5 dragons and renal tubular epithelium of 2/5 dragons from the initial mortality event. *Pasteurellaceae* bacteria of unknown were isolated from the lung and liver of each of the five dragons examined in 2014, and *Aeromonas hydrophila* was isolated in the lung of two of these animals: however, these were not considered contributory to pulmonary proliferation nor inclusion body formation. A summary of the signalment, gross and histological findings in each animal is summarized in Supplementary Table 1 and the histological lesions are shown in Figure 1.

**Figure 1.**
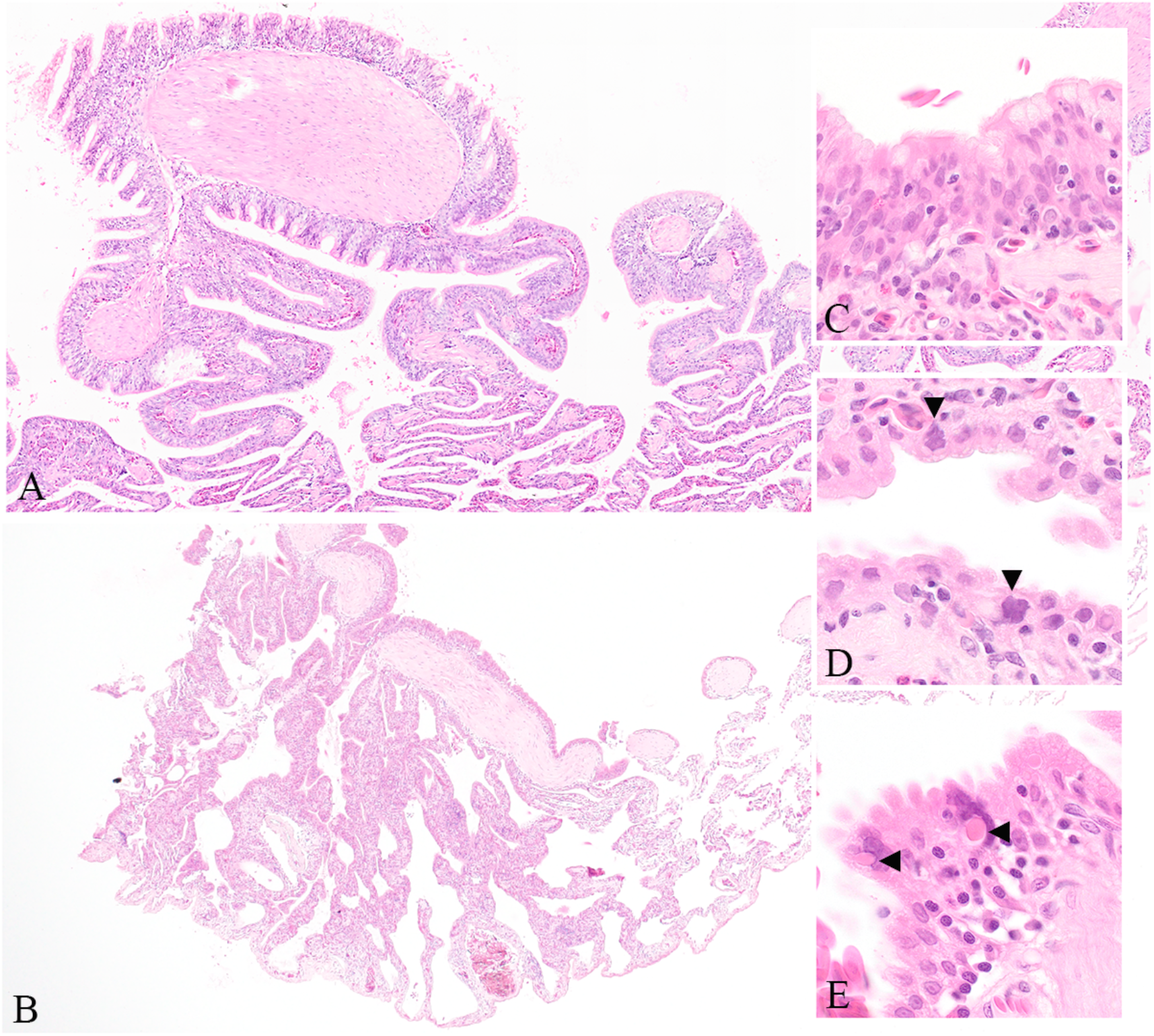
Photomicrographs of pulmonary lesions in central bearded dragons from the 2014 outbreak. Thickening of the normally squamous infundibular epithelium under low magnification (A) and the segmental nature of this change observed in an affected animal from the 2020 outbreak (B). Metaplasia of the normally squamous infundibular epithelium to pseudostratified columnar and multifocally ciliate (C) and multifocal epithelial cells with large, irregularly shaped, basophilic nuclei (arrowheads) (D). Bizarre atrial epithelial cells containing eosinophilic cytoplasmic inclusions (arrowheads) (E).

### PCR testing for pneumotropic viruses

All samples were PCR-negative for sunshineviruses, ferlaviruses and orthoreoviruses.

### Meta-transcriptomic pathogen discovery

Sequencing of a pooled rRNA-depleted RNA-seq library resulted in 81,188,754 raw reads. After filtering, 54,804,486 paired-end trimmed reads were generated, which were then *de novo* assembled into 266,118 contigs. Analyses of these read data revealed the presence of two DNA viruses, denoted here as bearded dragon circovirus (BDCV) and bearded dragon chaphamaparvovirus (BDchPV), with read abundances of 0.0006% and 0.003%, respectively. Virus-targeted PCR assays (Table 1) and Sanger sequencing were used to recover the full genome sequences of these two viruses. Virus-specific PCR was also used on spinal cord, lung, liver and kidney to provide insight into the distribution of these viruses. No other reptilian associated circoviruses and parvoviruses were detected using SRA screening.

### Genome characterization of a novel bearded dragon circovirus

We identified an abundant circovirus-like contig (1693 bp, 304 transcripts per million [TPM]) from our pooled lung, kidney and liver samples. The typical features of the genus *Circovirus* include a non-enveloped, icosahedral, single-stranded circular DNA (ssDNA) genome and size of approximately 1.5-2 kb. The full genome of BDCV was recovered through PCR assays and Sanger sequencing. The BDCV genome comprises 1,761 bp of circular DNA with a 43.8% GC content, encoding rep and cap genes of 308 and 218 amino acid residues, respectively.

The replicase protein of circoviruses introduces an endonucleolytic nick within the typical stem-loop structure with nonamer sequence 5'-AGTATTAC-3' in the intergenic region of the genome, thereby initiating the rolling circle replication (RCR). Comparison with other members of the *Circoviridae* revealed the presence of the classical conserved motifs, including RCR motifs, as well as SH3 helicase motifs (Table 2 and Figure 2).

**Table 2.**
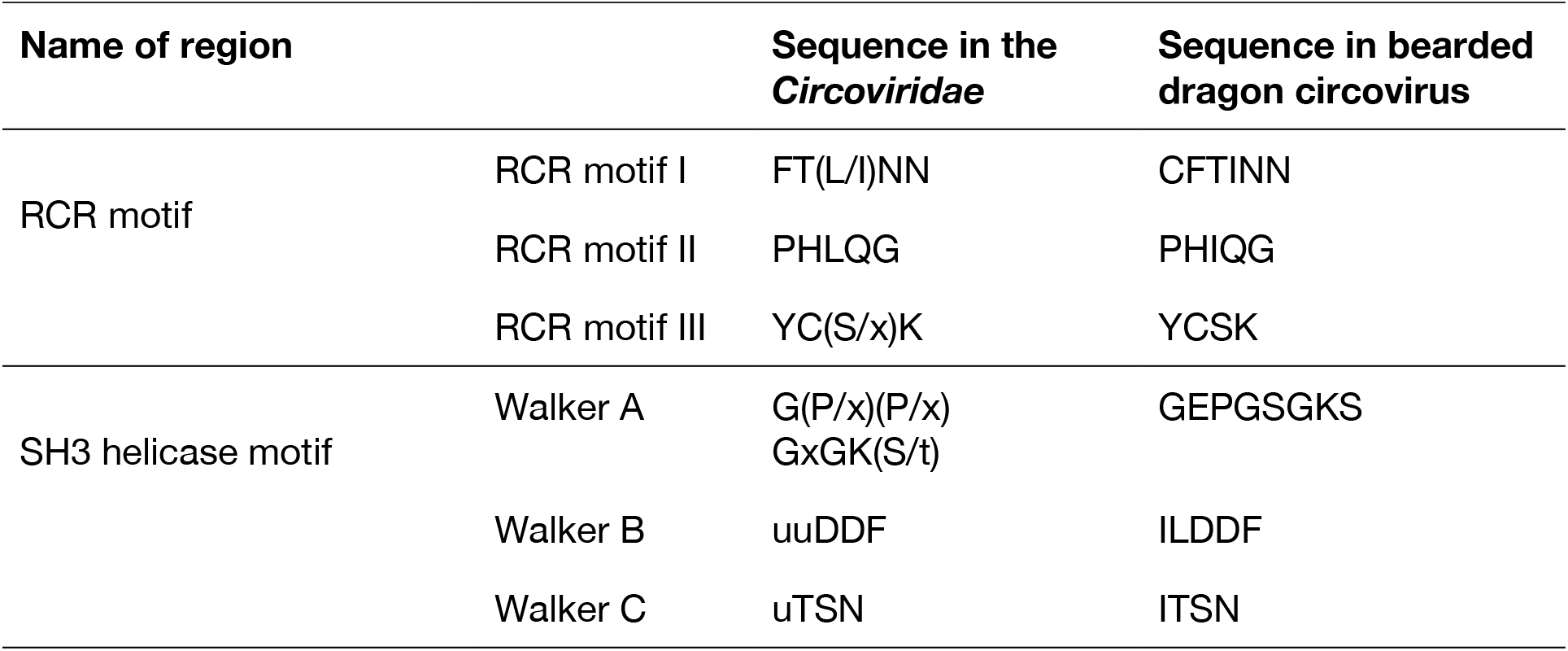
Rep protein (amino acid) motifs detected in the bearded dragon circovirus.

| Name of region |  | Sequence in the *Circoviridae* | Sequence in bearded dragon circovirus |
| --- | --- | --- | --- |
| RCR motif | RCR motif I | FT(L/I)NN | CFTINN |
|  | RCR motif II | PHLQG | PHIQG |
|  | RCR motif III | YC(S/x)K | YCSK |
| SH3 helicase motif | Walker A | G(P/x)(P/x)<br>GxGK(S/t) | GEPGSGKS |
|  | Walker B | uuDDF | ILDDF |
|  | Walker C | uTSN | ITSN |

**Figure 2.**
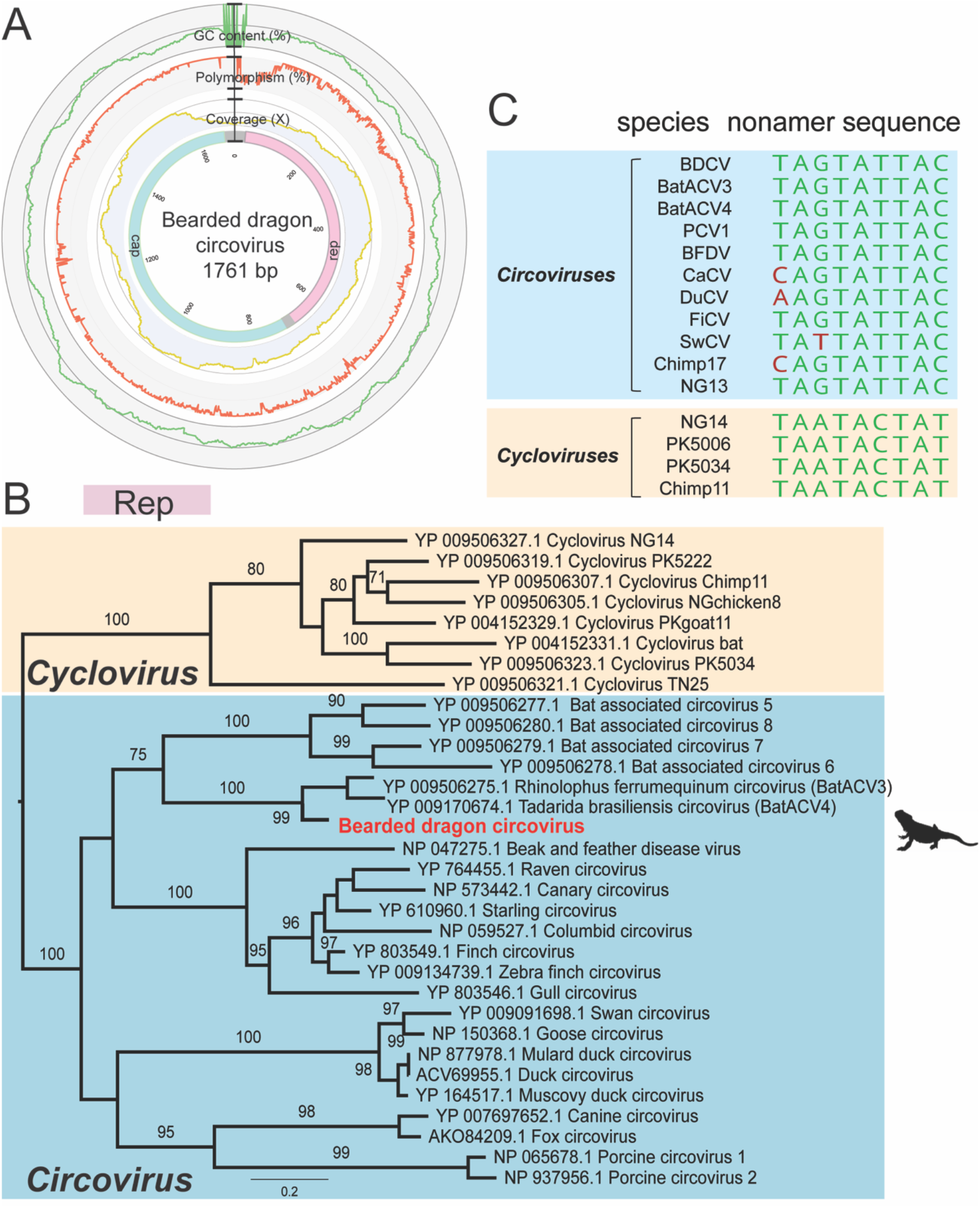
Genome characterization and phylogeny of bearded dragon circovirus. (A) Genome organization of bearded dragon circovirus. The outermost circles of the metadata ring represent the GC content (%, green), nucleotide polymorphism (%, orange) and read mapping coverage (yellow) of the genome. In the innermost circles, the proteins encoded by the replicase (Rep) and capsid protein (Cap) are labelled in pink and cyan, respectively. (B) Phylogenetic analysis of the Rep gene (replicase; DNA polymerase) of circoviruses, including members of the *Cyclovirus* and *Circovirus* genera. Bootstrap values > 70% are presented for key nodes (1,000 replicates).

The tree was midpoint rooted for clarity only. Scale bar shows the number of substitutions per site. (C) Comparison of nonamer sequences with closely related viruses in the *Circoviridae*.

To determine the evolutionary history of BDCV with respect to other circoviruses, we performed a phylogenetic analysis of the Rep protein. Notably, BDCV clustered robustly (99% bootstrap support; 77.85% amino acid pairwise identity in Rep genes) with a bat-associated virus lineage that includes *Tadarida brasiliensis* circovirus 1 (accession number YP_009170674.1) and *Rhinolophus ferrumequinum* circovirus 1 (YP_009506275.1), recently identified in metagenomic analyses of bat gut samples from Brazil and China, respectively.

### Identification of a highly divergent bearded dragon chaphamaparvovirus

We identified two abundant chaphamaparvovirus-like transcripts in the RNA-seq library. The *Parvoviridae* are a family of small, non-enveloped, dsDNA animal viruses with a linear genome of 4-6 kb in length. Chaphamaparvoviruses are a recently identified group of parvoviruses, likely representing a new genus, for a which a variety of component virus species have recently been discovered [33–36]. Four sets of bridging PCR assays were then designed to recover a near complete virus genome. This revealed that the virus genome comprised 4,181 nt with two distinct ORFs that encode the non-structural protein (NS, 633 aa) and the structural protein (VP, 600 aa). We further utilized specific primers to amplify the targeted NS (BDchPV_F2 and R1) and VP region (BDchPV_F4 and R3) for screening the virus distribution in different organs (Table 1).

Two conserved motifs were identified in the NS protein, including the putative endonuclease metal coordination motif ‘HLH’ and the helicase motif ‘GPASTGKS’ at positions 110-112 and 310-317, respectively (Figure 3). Additionally, the potential accessory proteins p10 (positions 74-358, 95 aa) and p15 (positions 136-543, 136 aa), previously identified in most other amniote-associated chaphamaparvoviruses, were both detected in BDchPV. The conserved PLA2 and G-rich motifs widely detected in other members of *Parvoviridae* were absent from the VP protein and hence similar to other chaphamaparvoviruses [chapparvoviruses] [37,38].

**Figure 3.**
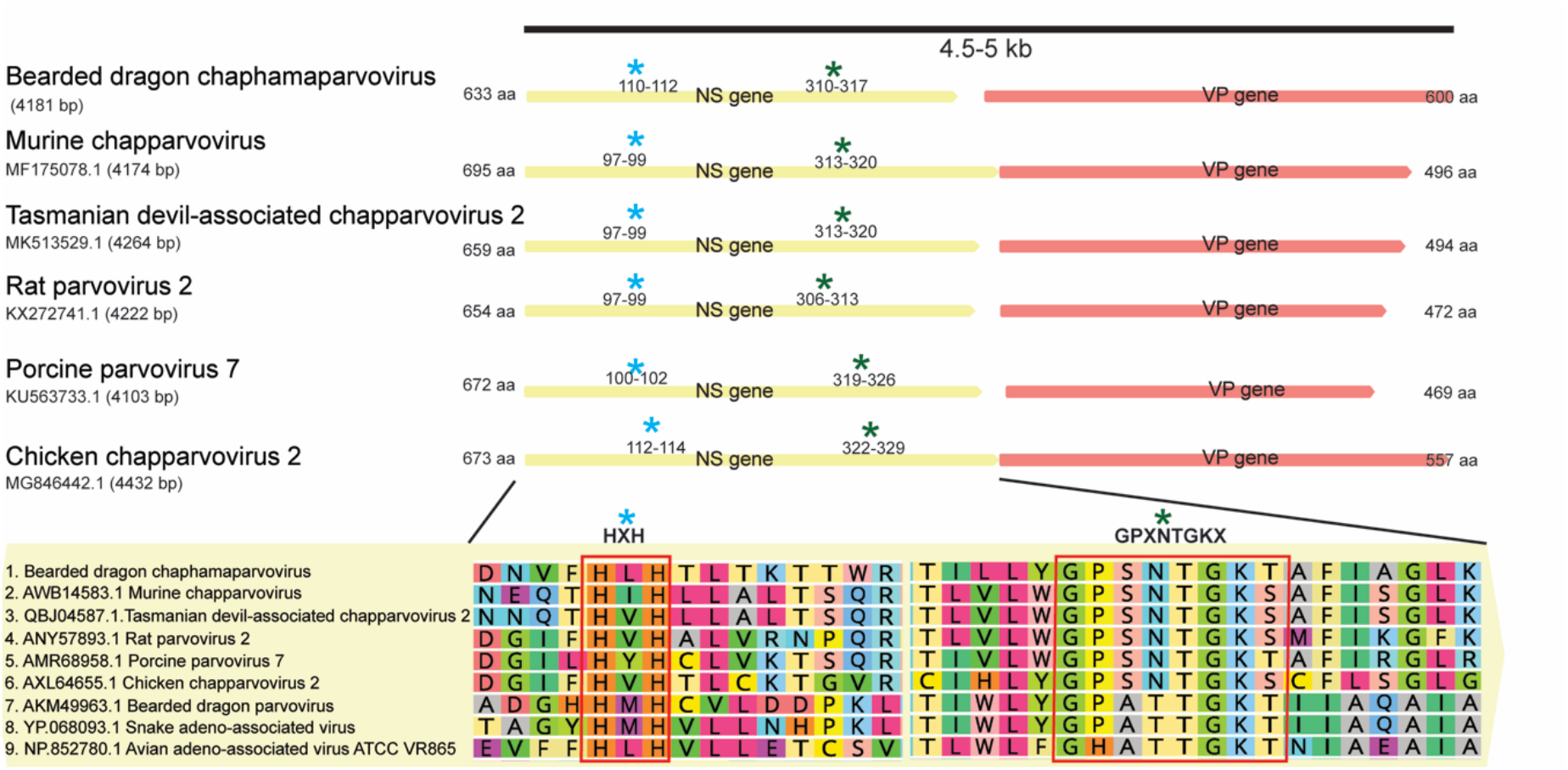
Bearded dragon chaphamaparvovirus in comparison to related chaphamaparvoviruses. The amino acid sequence size of the ORFs of each virus is shown. Yellow and orange boxes refer to the NS and VP genes, respectively. The classic motif features HXH (blue asterisk) and GPXNTGKX (green asterisk) on NS genes are labelled.

Phylogenetic analysis based on the predicted amino acid sequence of the complete NS protein revealed that BDchPV fell within the chaphamaparvovirus lineage, clustering with fish and crocodile-associated chaphamaparvoviruses. Tilapia parvovirus is a recently identified chaphamaparvovirus isolated from the feces of farmed tilapia and crocodiles in China [36]. However, the amino acid sequences of the novel BDchPV identified here shared only 42.1% and 41.2 % pairwise identity in NS gene and VP gene with this virus (accession no: QGX73405.1), respectively (Figure 4).

**Figure 4.**
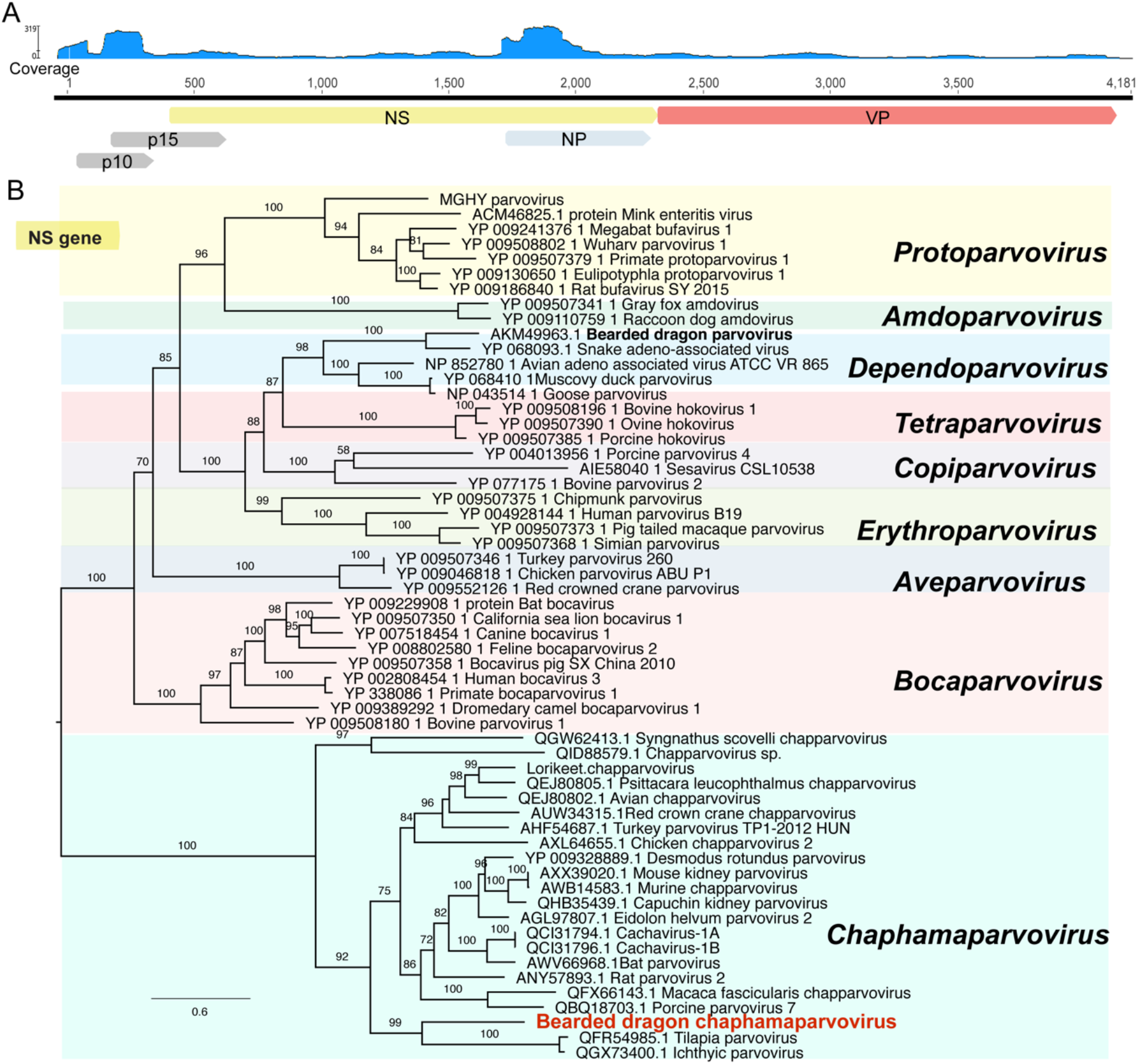
Genome characterization and phylogenetic relationships of central bearded dragon chaphamaparvovirus. (A) Genome depiction and reads mapping coverage of bearded dragon chaphamaparvovirus. (B) Phylogenetic analysis of the non-structural protein gene (NS gene) of parvoviruses, including other chaphamaparvoviruses and members of the *Dependovirus*, *Tetraparvovirus*, *Copiparvovirus*, *Erythroparvovirus*, *Protoparvovirus*, *Amdoparvoviru*s, *Aveparvovirus* and *Bocavirus* genera. Bootstrap values >70% were presented for key nodes (1,000 replicates). The tree was midpoint rooted for clarity only. Scale bar shows the number of substitutions per site.

### Prevalence of BDCV and BDchPV through PCR screening

The nucleic acid extracted from archived bearded dragon cases in the 2014 (n=5) and 2020 (n=12) outbreaks were screened using BDCV and BDchPV specific primers, respectively (Table 3). The circovirus was detected from kidney and lung samples from one animal (case: 10043.3) in the original respiratory disease outbreak. A total of 3/12 liver samples from the second outbreak were BDCV positive. In total, the chaphamaparvovirus was detected in the liver samples of 7/15 cases, including three original respiratory cases in 2014 and four additional cases in 2020 (Table 3). Sanger sequencing results from PCR products showed 95-99% nucleotide identity to the index cases.

**Table 3.**
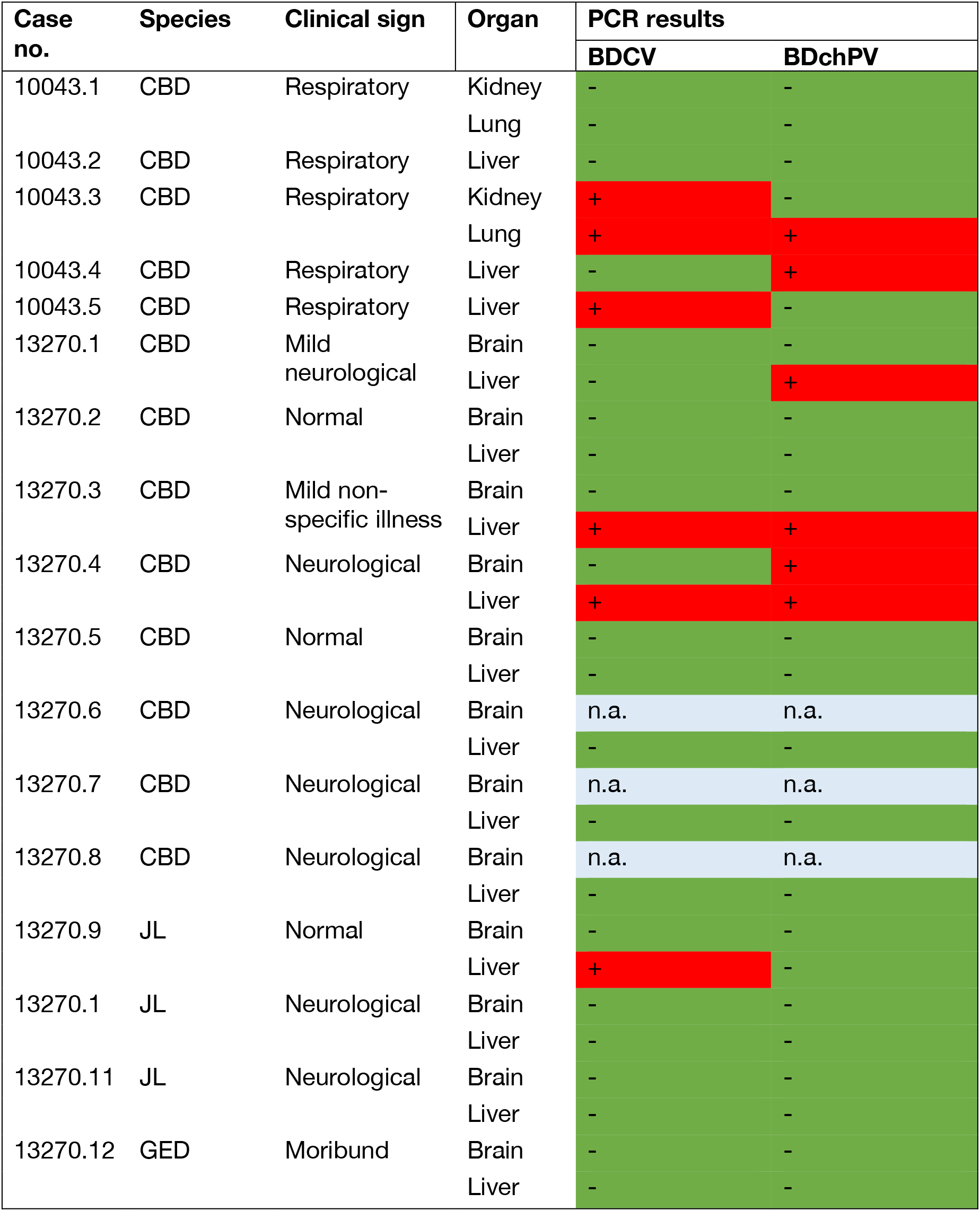
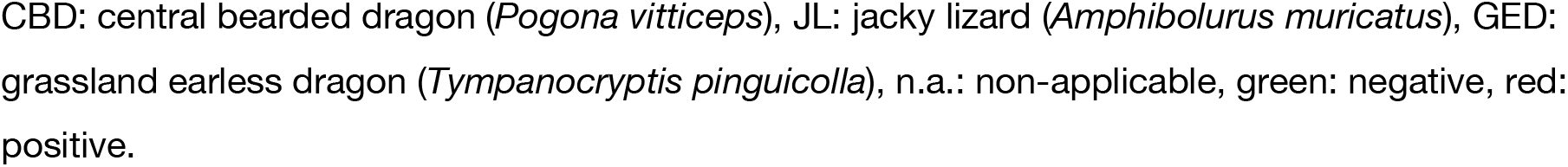
PCR testing of captive reptiles from the 2014 and 2020 outbreaks.

## Discussion

Our knowledge of the viruses of herpetofauna is expanding rapidly. Herein, we describe the meta-transcriptomic discovery and characterization of two complete genomes of DNA viruses, and their prevalence from mass mortality events of captive reptiles.

Circoviruses are one of the smallest DNA viruses that infect a wide range of vertebrates, with most pathogenic consequences in association with immunocompromised or immunosuppressed hosts, including canine [39], psittacine [40], porcine [41] and mink [42] species. In 2019, a circovirus was detected in the liver and gut of black-headed pythons (*Aspidites melanocephalus*) with spinal osteopathy [43]. The virus was then identified in tissues of a Boelen’s python (*Morelia boeleni*) and two annulated tree boas (*Corallus annulatus*). That investigation claimed to be the first to describe a reptilian circovirus, although other circo-like viruses had been described previously in testudines and squamates [44,45]. In all cases, the association between the circovirus and disease was either weak or absent. Endogenous circoviruses have also been detected in snake genomes [46]. It is notable that the bearded dragon circovirus newly identified here clustered with a bat-associated circoviruses lineage. However, these bat viruses were identified through metagenomic sequencing of environmental related samples (i.e. fecal viromes) such that their true hosts remain unclear.

Parvoviruses are associated with disease in a variety of host species, ranging from canines [49], livestock [50–53], rodents [36], and humans [54]. Reptile parvoviruses were identified in several snake species (snake adeno-associated virus) [11,55–59] , some lizard species [11] including bearded dragons [11,12]. To date, most of the exogenous reptilian parvoviruses identified belong to the genus *Dependoparvovirus* and it has been thought that these viruses require helper viruses (usually adenoviruses) for replication [13], although more recent work is challenging this notion [60]. Chaphamaparvoviruses (ChPVs) are a newly identified genus of parvoviruses [47]. Many ChPVs were previously unassigned members of the now-defunct *Chapparvovirus* genus; a genus that had been identified in rodents, birds, pigs, bats, Tasmanian devils, dogs, cats, primates and even invertebrates [33–37,48]. Although a ChPV was previously identified in crocodile feces [36], it is possible that this virus originated in the tilapia fish fed to these crocodiles. Our documentation of a novel and exogenous reptile chaphamaparvovirus in central bearded dragons extends the host range of these parvoviruses.

Interestingly, co-infection with circoviruses and parvoviruses is increasingly reported in different animal species, often in association with immunosuppression and exacerbated disease severity. For example, previous studies have indicated that Porcine circovirus 2 (PCV-2) associated diseases are augmented by concurrent viral infections such as Porcine parvovirus, and these viruses may serve as important cofactors in the pathogenesis of Porcine multisystemic wasting syndrome (PMWS) [61]. Similarly, Canine circovirus 1 (CaCV-1) and Canine parvovirus 2 (CPV-2) infection are linked to recurrent outbreaks of bloody diarrhea and sudden death in puppies [62], and a novel lorikeet chaphamaparvovirus co-infected with psittacine circovirus (beak and feather disease virus) in wild birds has been described [63]. Recently, a similar coexistence of Tasmanian devil-associated circovirus and chaphamaparvovirus was identified in a Tasmania devil metagenomic virome study, but with no disease association [64].

Despite our description of these pathogens, their roles in the disease syndromes described here remain uncertain and require additional investigation. Indeed, the examination of feeding and husbandry practices surrounding brumation is warranted, particularly since the animals that did not undergo brumation appeared to be unaffected in the mortality event. Although the association between the two viruses and proliferative lung disease was not clearly established, the role of these viruses as predisposing factors for disease or reduced fitness merits further investigation.

Reptiles comprise a considerable proportion of the vertebrate biomass of many ecosystems and play an important role in the food-web, and the delivery of ecosystem services through pollination, seed dispersal and pest control. Australia has the highest diversity of lizards of any country, yet our understanding of the biological threats to these animals is scarce. It has been difficult to obtain an accurate understanding of the health status of wildlife that are free-living, maintained for research, education, or species recovery, using traditional diagnostic techniques. This study illustrates the value of metagenomic approaches to disease investigation as a complement to traditional histology and pathogen culture. The identification of bearded dragon circovirus and the bearded dragon chaphamaparvovirus provides insight into reptilian viral diversity and reptile health, and undoubtedly merits broader surveillance.

## Data Availability

The genome sequence of BDCV and BDchPV are available on NCBI/GenBank (accession numbers: MT732118 and MT73219). Raw sequencing reads are available at the Sequence Read Archive (SRA) under accession PRJNA644669.

## Author Contributions

Conceptualization, E.C.H., K.R. and W.-S.C.; methodology, W.-S.C., C.-X.L., T.H.H., J.H. and K.R.; software, W.-S.C.; laboratory analysis, W.-S.C., C.-X.L., J.-S.E. and K.R.; formal analysis, W.-S.C.; investigation, W.-S.C.; resources, K.R. and E.C.H.; data curation, W.-S.C.; writing—original draft preparation, W.-S.C. and K.R.; writing—review and editing, W.-S.C., C.-X.L, J.H., J.-S.E., T.H.H., E.C.H. and K.R.; visualization, W.-S.C. and K.R.; supervision, E.C.H., K.R. and J.-S.E.; funding acquisition, E.C.H., K.R. All authors have read and agreed to the published version of the manuscript.

## Acknowledgments

This research was supported by Taronga Conservation Society Australia. We thank Wendy Ruscoe, Jacqui Richardson and Professor Arthur Georges, University of Canberra for their contribution of specimens, history and context. Paul Thompson, Taronga Wildlife hospital is acknowledged for his role in microbial culture. The authors thank CSIRO’s Australian Centre for Disease Preparedness and Dr. Mark Crane for contributing the viral culture results and acknowledge The University of Sydney HPC service (Artemis) for providing high-performance computing resources that have contributed to the research reported in this paper. We also thank Dr. Wen-Lin Wang for her advice and support.

