## Supplementary Table 1 for "Meta-transcriptomic discovery of a divergent circovirus and a chaphamaparvovirus in captive reptiles with proliferative respiratory syndrome"

**Table S1**. Presentation, pathology in the central bearded dragon morbidity and mortality event, 2014.

| Case ID | Signal ment | Gross lesions | Proliferative respiratory Epithelium | Pneum onia I/N | Eosinophilic Cytoplasmic IB |
|---|---|---|---|---|---|
| 10043.1 | Male/Adult | Miliary white hepatic foci. Pleural petechiae, pulmonary congestion, mucoid yellow exudate in mesobronchus. Small testes. Fair body condition. | 4D | 4D/2F | Lung, kidney, liver |
| 10043.2 | Female/Adult | Mottled hepatic parenchyma. Pulmonary congestion. Multifocal haemorrhagic ova. Fair body condition. | 1D | 2MF/2MF | Liver |
| 10043.3 | Female/Adult | Mulitofocal haemorrhagic ova. Good body condition | 1D | 3D/0 | 0 |
| 10043.4 | Male/Adult | Dehydrated. Hepatic atrophy. Blood surrounding bolus of food in large intestine. Small testes. Poor body condition. | 4D Binucleate cells and syncytia | 1D/1D | Lung, kidney, liver |
| 10043.5 | Female/Adult | Mottled hepatic parenchyma. Pulmonary congestion and unilateral mesobronchial blood clot. Small ova. Poor body condition. | 3D | 3D/3D | Lung, nerve cell body |

Proliferation of respiratory epithelium graded on a scale of 0-4 in ascending severity. Inflammation (I) graded on a scale of 0-4 in ascending severity. Necrosis (N) graded on a scale of 0-4 in ascending severity. D – diffuse, S – segmental, MF – multifocal, F – focal tracts. IB: inclusion bodies.
